# Biochemical alterations precede neurobehavioral deficits in a novel mouse model of Friedreich ataxia

**DOI:** 10.1101/2021.04.05.438486

**Authors:** Marta Medina-Carbonero, Arabela Sanz-Alcázar, Elena Britti, Fabien Delaspre, Elisa Cabiscol, Joaquim Ros, Jordi Tamarit

**Affiliations:** Dept. Ciències Mèdiques Bàsiques, Fac. Medicina, IRBLleida, Universitat de Lleida. Lleida.

**Keywords:** Friedreich Ataxia, Iron-sulfur, Mitochondria, Oxidative stress, OXPHOS

## Abstract

Friedreich Ataxia (FA) is a rare neuro-cardiodegenerative disease, caused by partial deficiency of frataxin, a mitochondrial protein. This deficiency is caused by the presence of a GAA triplet expansion in the first intron of the frataxin gene or, in some patients, by point mutations. Generating mouse models mimicking FA has been challenging, as this disease is manifested when frataxin levels are below a pathological threshold. In the present work, we have characterized a new mouse model of FA (FXN^I151F^) based on a pathological point mutation (I154F) present in some FA patients. These mice present very low frataxin levels in all tissues and display neurological deficits resembling those observed in FA patients. We have also observed decreased content of components from OXPHOS complexes I and II, decreased aconitase activity, and alterations in the antioxidant defenses. Remarkably, these biochemical alterations precede the appearance of neurological symptoms and present a different profile in heart and brain or cerebellum. The FXN^I151F^ mouse is an excellent tool for analyzing the consequences of frataxin deficiency in different tissues and for testing new therapies.

## INTRODUCTION

Friedreich Ataxia (FA) is a rare, inherited recessive disease first described by Nikolaus Friedreich, a German pathologist, in 1863. FA is characterized by progressive gait and limb ataxia with associated limb muscle weakness, absent lower limb reflexes, extensor plantar responses, dysarthria, and decreased vibratory sense and proprioception (Parkinson *et al*, 2013). Most FA patients also present hypertrophic cardiomyopathy, being cardiac dysfunction the leading cause of death (Tsou *et al*, 2011). The disease is caused by mutations in a gene on chromosome 9, called FXN (for frataxin) or X25 that result in decreased frataxin content or expression. In most patients, frataxin deficiency is caused by the presence of a GAA triplets expansion in the first intron of FXN gene. This expansion is found in both alleles and compromises frataxin expression (Campuzano *et al*, 1996). Around 4% of patients are compound heterozygous for a GAA expansion and a FXN point mutation or deletion (Galea *et al*, 2016).

Decreased expression of frataxin is associated with mitochondrial dysfunction, iron and calcium unbalance, and increased oxidative stress. The function of frataxin and the mechanisms causing such cellular disturbances are not completely understood. It has been shown that frataxin localizes to mitochondria where it regulates the activity of cysteine desulfurase, an enzyme required for the biosynthesis of iron-sulfur clusters (Gervason *et al*, 2019). Therefore, it is generally accepted that frataxin activates iron-sulfur biogenesis in eukaryotes. Nevertheless, iron-sulfur deficiency is not a universal consequence of frataxin deficiency, suggesting that the role of frataxin in iron-sulfur biogenesis is not essential (Alsina *et al*, 2018a). Frataxin deficiency also causes iron overload, and different mechanisms have been proposed to explain this fact. First, compromised iron-sulfur biogenesis could interfere with iron sensing by Iron Regulatory Proteins, leading to constitutive activation of iron uptake (Seznec *et al*, 2005)(Li *et al*, 2008)(Whitnall *et al*, 2008). Second, frataxin binds iron and presents ferroxidase activity. Therefore, it could be involved in iron chelation and/or detoxification inside mitochondria (Gakh *et al*, 2006). Futhermore, large oligomers of frataxin are formed *in vitro* in the presence of iron that could be involved in iron storage. Nevertheless, the *in vivo* relevance of such structures is controversial (reviewed in (Tamarit *et al*, 2021)). Third, it has also been described that an extramitochondrial form of frataxin would bind and regulate Iron Regulatory Protein 1 (Condò *et al*, 2010), although the existence of such extramitochondrial form of frataxin is a matter of debate.

Besides iron and/or iron-sulfur biogenesis, other processes are altered in frataxin-deficient cells. Several observations suggest that oxidative stress could play a central role in FA. Hypersensitivity to oxidative damage has been observed in FA models and patients, and it has been linked to deficient activation of the NRF2 pathway in response to oxidative insults. This pathway is required to induce the expression of antioxidant defenses (Paupe *et al*, 2009)(Shan *et al*, 2013). The central role of oxidative stress in FA is also highlighted by recent reports indicating that frataxin-deficient cells are hypersensitive to erastin, a drug which causes GSH depletion and which is a well-known inducer of ferroptosis (Cotticelli *et al*, 2019)(Du *et al*, 2020). Ferroptosis is a form of regulated cell death, which is triggered by the accumulation of lipid peroxidation products in membranes. Frataxin deficiency has also been shown to impact on different pathways related to mitochondrial function. These include the OXPHOS system (Tamarit *et al*, 2016) (Lin *et al*, 2017), mitochondrial calcium efflux (Britti *et al*, 2021), mitochondrial permeability pore opening (Purroy *et al*, 2018), pyruvate dehydrogenase complex (Purroy *et al*, 2020), and endoplasmic reticulum-mitochondria contacts (Rodríguez *et al*, 2020), among others

Generating mouse models to recreate FA has been challenging, as this disease is manifested when frataxin levels are below a pathological threshold. Therefore, animals must express some frataxin to be viable, but such expression may be lower than the pathological threshold required to trigger the disease. In this regard, complete FXN Knock-out (K.O.) mouse are embryonically lethal (Cossée *et al*, 2000), while a mice model presenting a 70% reduction in frataxin content (known as the KIKO model) displays a mild phenotype (McMackin *et al*, 2017). Several “humanized” models have been developed in which a human FXN gene carrying GAA expansions is used to rescue the KO mice. These models have a very mild phenotype that is only observed in aged animals, limiting its use to test new therapies (Al-Mahdawi *et al*, 2006). Conditional tissue-specific KO mice have also been developed (Puccio *et al*, 2001). These models can be used to analyse the specific consequences of frataxin deficiency in a specific tissue, but do not mimic the biochemical problem encountered by patients (the presence of limited amounts of frataxin in all tissues). Therefore, in the present work, we introduce a new model of FA based on the I154F pathological point mutation. This mutation was first reported in 3 unrelated Italian FA patients which were compound heterozygotes for the GAA triplet-repeat expansion and the I154F point mutation. The clinical phenotype of these patients was indistinguishable from that of individuals homozygous for the GAA expansion (Filla *et al*, 1996). Analysis in cultured mammalian cells revealed that the I154F mutation compromised the solubility of frataxin intermediate form resulting in lower content of mature frataxin (Li *et al*, 2013). In the present work we show that homozygous mice for the equivalent mutation (I151F) present very low frataxin levels, neurological deficits and biochemical alterations which mimic those observed in patients with FA. Therefore, this mouse model is an excellent tool for analyzing the consequences of frataxin deficiency and for testing new therapies.

## RESULTS

### Generation of frataxin I151F knock-in mice

FXN^I151F/wt^ heterozygous mice (C57BL/6J-Fxn^em10(T146T,I151F)Lutzy/J^) were obtained from the Jackson Laboratory (Stock Number 31922). To generate these mice, CRISPR/Cas9 was used to introduce the ATC→TTC mutation in the I151 codon from murine *Fxn.* The resulting I151F substitution is equivalent to the human pathogenic I154F missense mutation (Figure 1A). An additional silent mutation is present in the T146 codon (ACC→ACT). This change was deliberately co-introduced with the desired mutation in order to destroy the guide dependent PAM recognition site. Inter-crossing of these mice resulted in progeny of all possible genotypes. From 180 mice born, 48 (26,7%) were WT mice (FXN^wt/wt^ homozygous for wild type FXN), 91 (50,5%) were HET mice (FXN^I151F/wt^, heterozygous for wild type FXN and FxnI151F), and 41 (22,8%) were Fxn^I151F^ mice (FXN^I151F/I151F^, homozygous for the FxnI151F allele) (Figure 1D). Therefore, progeny was in the expected mendelian ratios indicating that MUT mice were viable.

**Figure 1.**
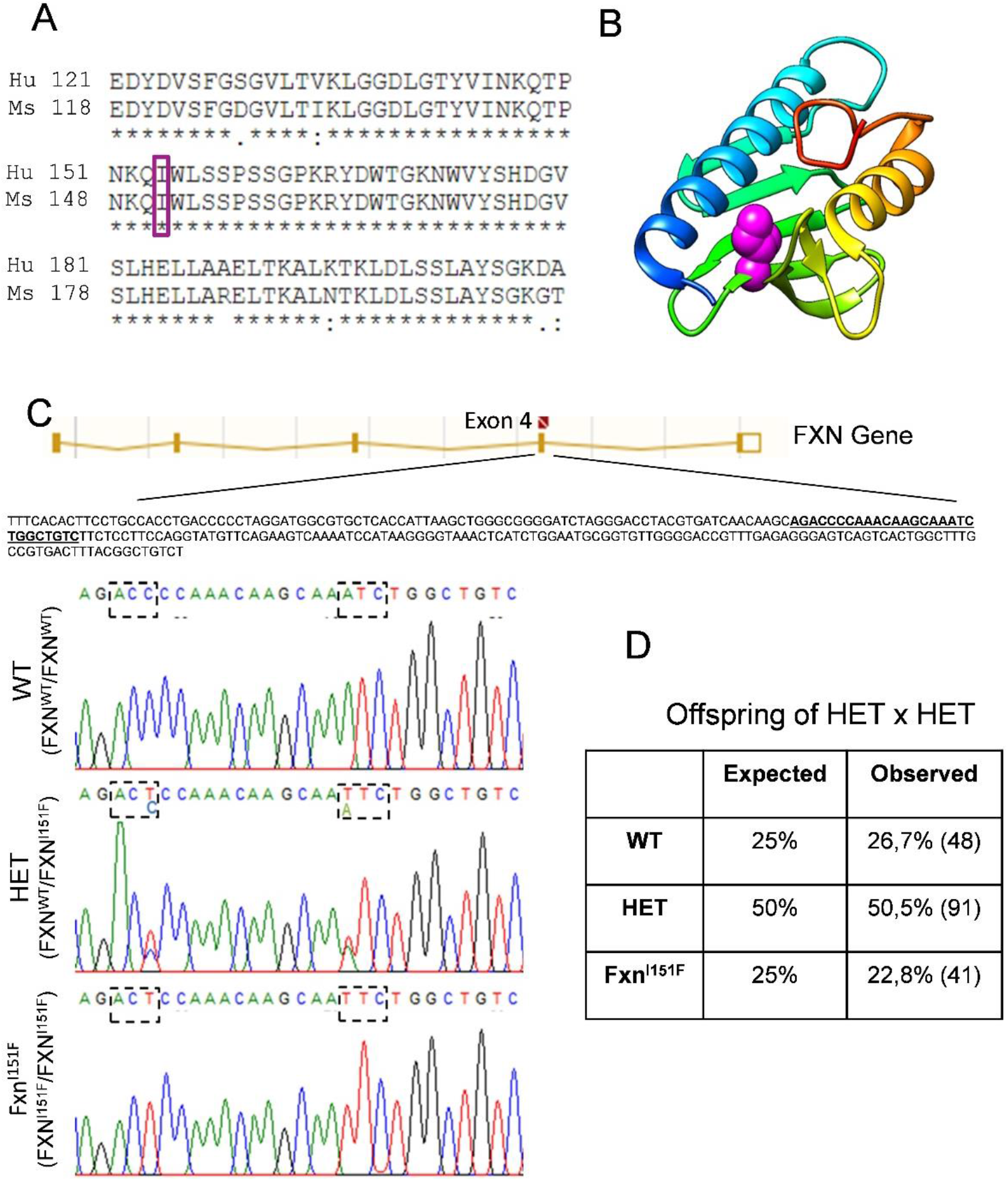
Generation of the FxnI151F model. A, alignment of the C-terminal region from human (Hu, Uniprot Q16595) and mouse (Ms, Uniprot O35943) frataxin protein sequences. Amino acid I151 in mice corresponds to the I154 in humans (boxed). B, ribbon representation of human frataxin (pdb code 3s4m) showing the position of I154 (spacefill, in violet). Ribbons are colored according to sequence, from dark blue (N-terminal) to red (C-terminal). Molecular graphics were performed with the UCSF Chimera package. C, DNA sequence analyses of WT, HET and Fxn^I151F^ mice. The top panel shows a schematic diagram of FXN Gene. The amplified sequence used in genotyping assays includes exon 4 and is indicated. The region containing the mutations is highlighted and shown below. Dashed boxes indicate the ACC→ACT (silent) and ATC→TTC (I151F) mutations introduced by gene editing in the frataxin gene. D, genotyping of litters from HET intercrosses.

### Frataxin protein levels are markedly reduced in Fxn^I151F^ mice

To determine the effect of I151F mutation in frataxin content, we measured the levels of this protein in brain, cerebellum and heart from 21 and 39-week old WT, HET and Fxn^I151F^ mice. By western blot we observed that the content of mature frataxin in HET mice was approximately 50% of that observed in WT mice, while a residual 3% mature frataxin content was observed in Fxn^I151F^ mice (Figure 2A). We also analyzed the potential presence of an insoluble frataxin intermediate form, which was reported in cells overexpressing human I154F frataxin (Li *et al*, 2013). With this purpose, we overexposed western blot membranes in order to analyze the presence of intermediate or high molecular weight forms of frataxin. Tissue homogenates were prepared in the presence of 4% SDS, a concentration high enough to solubilize the previously reported I154F insoluble frataxin intermediate form. We could detect the presence of some high molecular weight bands in WT homogenates, which were more clearly observed in heart. One of these bands was close to the expected mobility for mouse full-length frataxin (22.9 kDa) (Figure 2B). Nevertheless, these high molecular weight forms represented less than 1% of the chemiluminescent signal from the mature form, and were also decreased in homogenates from HET and Fxn^I151F^ mice. Therefore, we can exclude the presence of an insoluble intermediate frataxin proteoform in mice harboring the I151F mutation. We also analyzed the content of frataxin in homogenates from spinal cord, dorsal root ganglia (DRG), liver, pancreas and skeletal muscle. Gels were overloaded in order to detect residual frataxin levels in Fxn^I151F^ mice. This analysis confirmed that the I151F mutation causes a marked loss of frataxin content in all mouse tissues (Figure 2C).

**Figure 2.**
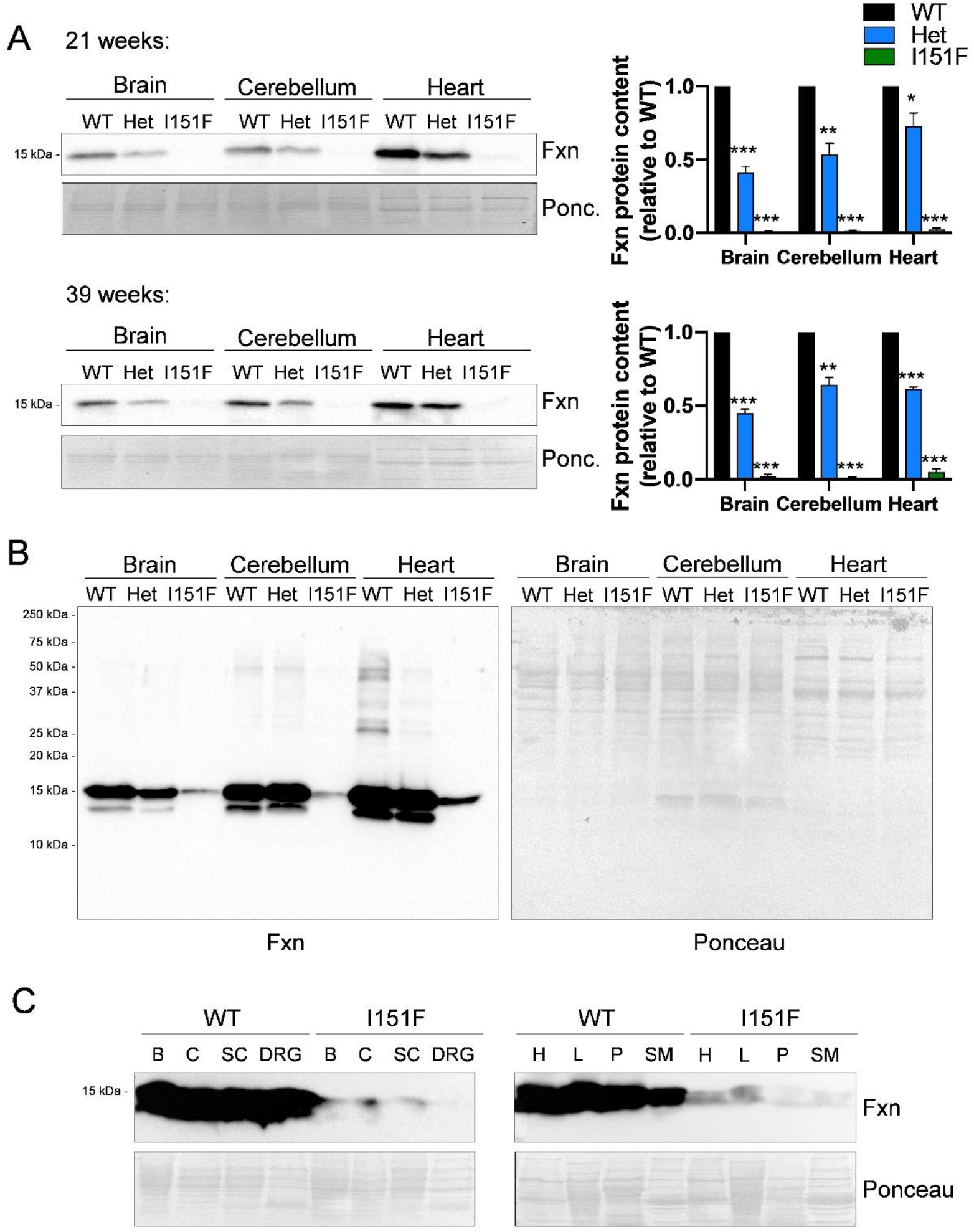
Fxn^I151F^ mice present decreased frataxin content. A, relative mature frataxin (15kDa) content analysed by western blot in brain, cerebellum and heart homogenates from WT, HET and Fxn^I151F^ mice. B, a representative image from an overexposed frataxin western blot membrane loaded with homogenates from 39-week old mice. Whole membrane is shown to indicate the absence of insoluble full-lenght or frataxin intermediate proteoforms. Please note that mature frataxin (15kDa) signal from WT and HET homogenates is saturated. C, representative image from frataxin western blot loaded with 50 μg from WT and Fxn^I151F^ homogenates from brain (B), cerebellum (C), spinal cord (SC), dorsal-root ganglia (DRG), heart (H), liver (L), Pancreas (P), and skeletal muscle (SM) from 39-week old mice. Membranes were overexposed to allow visualization of mature frataxin in homogenates from Fxn^I151F^ mice.

### Fxn^I151F^ mice present decreased weight gain

Body weight of WT and Fxn^I151F^ mice were measured from birth every two weeks. Weight gain was similar in WT and Fxn^I151F^ mice until 10 weeks of age. From that age on, weight gain was lower in Fxn^I151F^ mice than in WT mice. Therefore, significant differences in weight were observed between WT and Fxn^I151F^ mice from 16 week of age onward (Figure 3A and B). The relative difference in weight between WT and Fxn^I151F^ mice was progressive and was similar in both females and males. At 39 weeks of age, Fxn^I151F^ mice presented on average a 23% decrease in weight, when compared with WT mice (Figure 3C and D).

**Figure 3.**
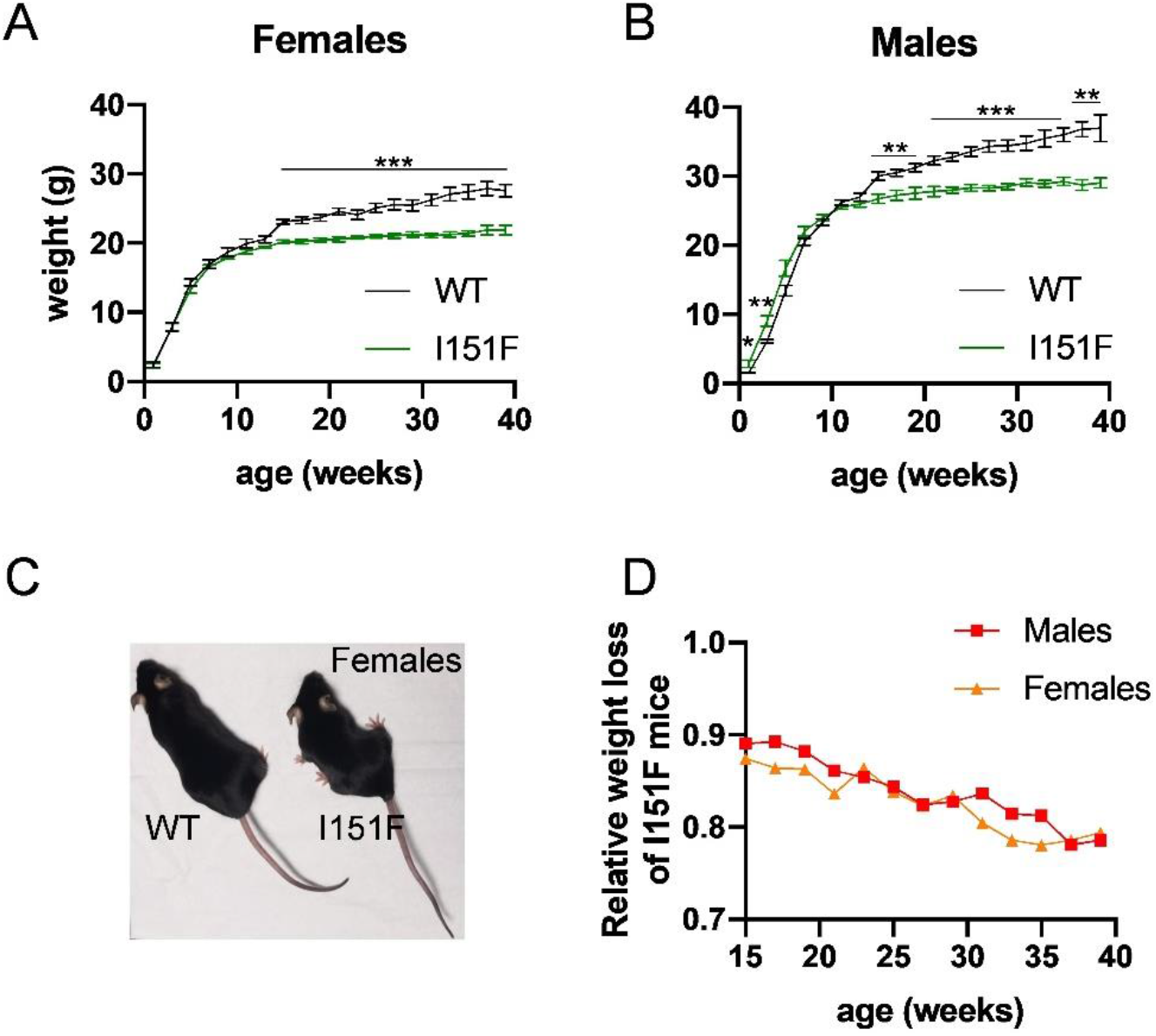
Fxn^I151F^ mice present decreased weight gain. A and B, body weight of WT and Fxn^I151F^ mice up to 39 weeks of age. C, representative image from WT and Fxn^I151F^ female mice. D, average weight loss of Fxn^I151F^ mice relative to WT mice of the same age.

### Fxn^I151F^ mice exhibit neurological deficits

To determine whether frataxin deficiency caused by the I151F mutation impacted the behavior of Fxn^I151F^ mice, we subjected these mice to several motor behavioral tasks. WT mice were also analyzed as a control. The number of mice used for each analysis is summarized in supplemental table 1. The motor coordination ability was assessed on a rotarod treadmill. This analysis was performed every two weeks, from 15 to 39 weeks of age. As shown in figure 4A, Fxn^I151F^ mice showed decreased coordination ability when compared with WT. Differences were statistically significant from week 21 onward. Forelimb strength was assessed using a hang-wire test, in which mice were assessed for their ability to hang on to a horizontal wire by their forepaws. Analysis were performed every six weeks (starting at week 21). Fxn^I151F^ mice fell off the wire quicker than WT mice. Statistically significant differences were obtained for 27, 33 and 39 week-old mice (Figure 4B). Locomotor activity tests were performed using an open-field beam-breaker activity monitor. These analyses were performed every 6 weeks (starting at week 21) up. Fxn^I151F^ mice exhibited significantly reduced average velocity, ambulatory distance (total distance covered by the mice within a specific time), and number of crossings than WT mice (Figure 4C). Finally, we assessed gait ataxia using paw print analysis every 6 weeks (starting week 21). Significant differences between Fxn^I151F^ and WT mice were detected in 39-week old mice, but not previously. At this age, Fxn^I151F^ mice displayed reduced hind and front limb stride length when compared with WT. This suggests that Fxn^I151F^ mice present a progressive ataxic gait (Figure 4D). Overall, the results presented in this section indicate that Fxn^I151F^ mice present progressive neurologic defects that are not observed before 22 weeks of age.

**Figure 4.**
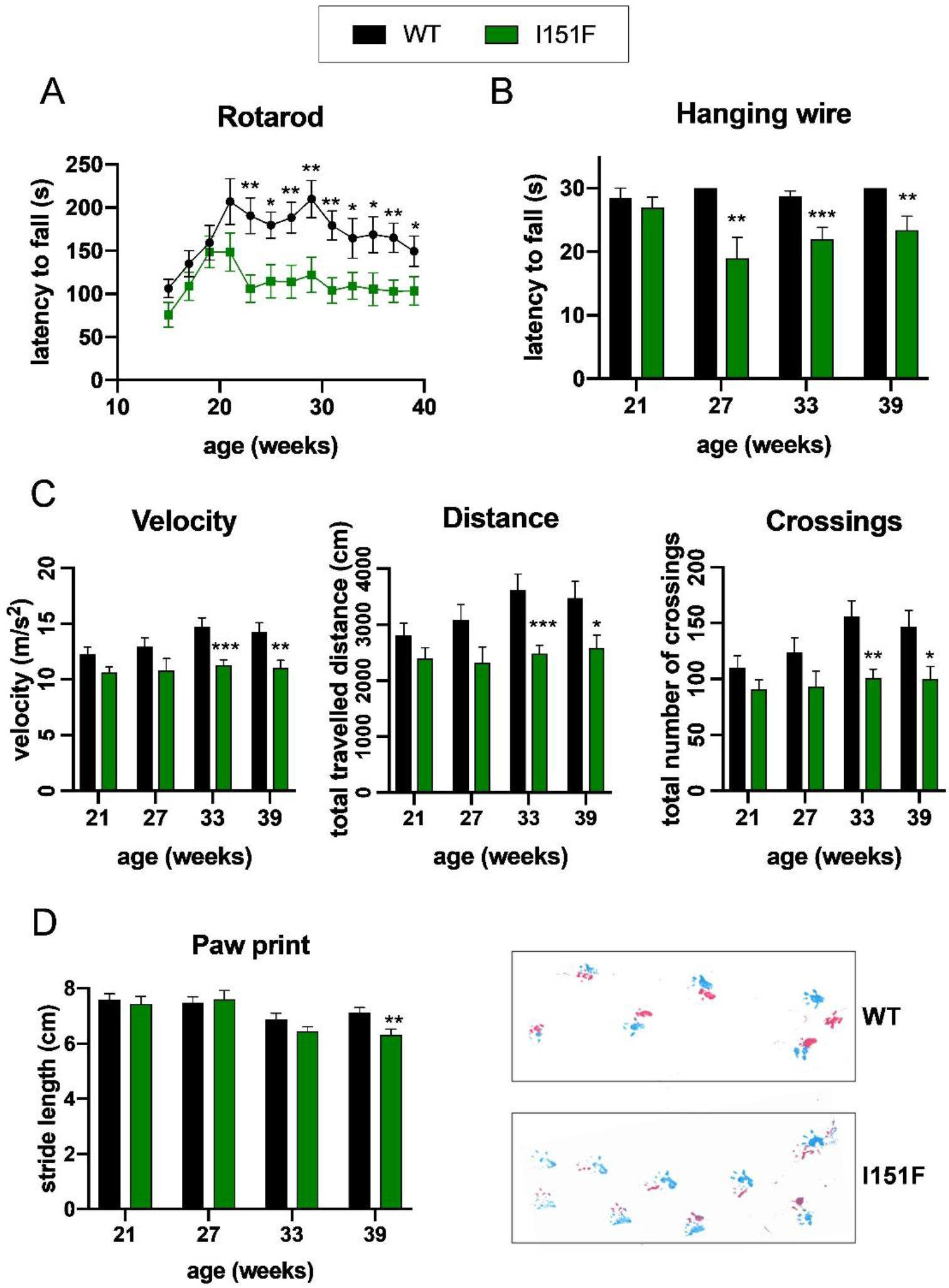
Neurological deficits in Fxn^I151F^ mice. Rotarod, Open field, hang wire and Paw print analysis in WT and Fxn^I151F^ animals. A, rotarod test in mice up to 39 weeks of age. Fxn^I151F^ animals presented a decreased latency to fall from 21 weeks onward. B, hanging-wire test in mice up to 39 weeks of age. Fxn^I151F^ animals presented a decreased latency to fall from 27 weeks onward. C, open field test. Fxn^I151F^ mice showed significant decline in velocity, total distance traveled and number of crossings from 33-weeks onward. D, gait footprint analysis revealed abnormalities in walking patterns in Fxn^I151F^ mice, which displayed significantly reduced stride length at 39-weeks of age. Representative walking footprint patterns are shown, in which pink and blue marks represent hind and fore paws respectively. Values are shown as mean ± SEM. Number of animals used in each analysis is indicated in supplemental table 1.

### Analysis of frataxin-related proteins by targeted proteomics

We were interested in analyzing the biochemical consequences of frataxin deficiency in nervous and cardiac tissues before (21-weeks) and after the onset of neurological defects (39-weeks). With this purpose, we decided to focus on several proteins or pathways that have been related with frataxin. In this regard, it has been described that frataxin deficiency causes loss of iron-sulfur containing proteins (Rotig *et al*, 1997), decreased function of the OXPHOS system (Lin *et al*, 2017), changes in the content of superoxide dismutases (Irazusta *et al*, 2006), and decreased content of the pyruvate dehydrogenase component PDH A1 (Purroy *et al*, 2020). Also, frataxin has been reported to interact with components of the iron-sulfur biosynthesis machinery (Patraa & Barondeaua, 2019), the OXPHOS system (Gonzalez-Cabo *et al*, 2005), and with the mitochondrial chaperone GRP75 (Shan *et al*, 2007) (Dong *et al*, 2019). Therefore, in order to explore these pathways in Fxn^I151F^ mice, we set up a targeted proteomics method to analyze the content of proteins related with these pathways in tissues collected from WT and Fxn^I151F^ mice. We used an SRM-based targeted proteomics approach in which proteotypic peptides from each protein were detected in a LC-triple quadrupole mass spectrometer. This approach allows the quantitation of several proteins in multiple samples with high reproducibility. The proteins analyzed are listed in table 1. They were two components of the tricarboxylic acids cycle (CS and Aco2), several components of the OXPHOS system (SDHA, SDHB, CY1, QCR2, COX2, ATPA, and ATPB), two mitochondrial chaperones (HSP60 and GRP75), three components of the Pyruvate dehydrogenase complex (PDHA1, DLAT and DLDH), and two superoxide dismutases (SOD1 and SOD2). Three glycolytic enzymes where also included in the analysis (GAPDH, PKM, and Enolases A, B, G) in order to analyze a potential unbalance between respiratory and glycolytic pathways and/or between mitochondrial and cytosolic pathways. Two of the proteins analyzed contain iron-sulfur clusters (ACO2 and SDHB), while two of them (CYC and CY1) contain heme groups. The relative content of these 21 proteins was analyzed in brain, cerebellum and heart from WT and Fxn^I151F^ mice, sacrificed at 21 or 39 weeks of age (in the case of enolases, ENOG only in nervous systems and ENOB only in heart). The mean CV of the measurements between replicas was <10%, indicating a high reproducibility of the data obtained. Results are shown in figure 5A. Histograms indicate that significant differences in the content of several proteins were observed between WT and Fxn^I151F^ mice. The net representation shown in figure 5B, allows a comparison of the relative content of the proteins (ratio Fxn^I151F^ /WT) at 21 and 39 weeks of age. It can be appreciated that the proteins more affected were ACO2 and the two components of OXPHOS complex II (SDHA and SDHB). These proteins showed a marked decrease in cerebellum and brain from Fxn^I151F^ mice, both at 21 and 39 weeks. In heart, we could only observe loss of complex II in 21 weeks-old mice, but not in 39-week old mice, while ACO2 content was not altered in heart at any age. Some other components of the OXPHOS system showed minor changes: the complex III components QCR2 or CY1 were found decreased either in brain (21-weeks), cerebellum (21 and 39 weeks) or in heart (21 weeks); ATPA and ATB were found increased in 39-week old cerebellum and heart. A second group of proteins showing changes in their content were the antioxidant enzymes SOD1 and SOD2. In heart, both enzymes were induced in 21-week old mice, while their levels decreased in 39-week old mice. Some induction of SOD2 was also observed in brain at 21 and 39 weeks. Regarding PDH components, induction of both PDHA1 and DLAT was found in 39-week old heart. In contrast, DLDH (the E3-component of PDH complex) was found decreased in 39-week old heart and cerebellum. No major changes were observed in the content of the mitochondrial chaperones analyzed, nor in the content of glycolytic enzymes. Overall, these results indicate that frataxin deficiency does not cause a general loss of mitochondrial proteins. Instead, it causes specific changes in the content of certain proteins. Also, the net representation indicates how those changes evolve as mice age. In this regard, brain and cerebellum do not experience major differences in the Fxn^I151F^ /WT protein ratio between 21-week old and 39-week old mice. Therefore, biochemical alterations precede the neurological defects described in the previous section. Instead, the heart shows differences in the Fxn^I151F^ /WT protein ratio between 21 and 39 week old mice, suggesting that the consequences of frataxin loss in this tissue are progressive. Those proteins presenting changes in their content were further explored.

**Table 1.**
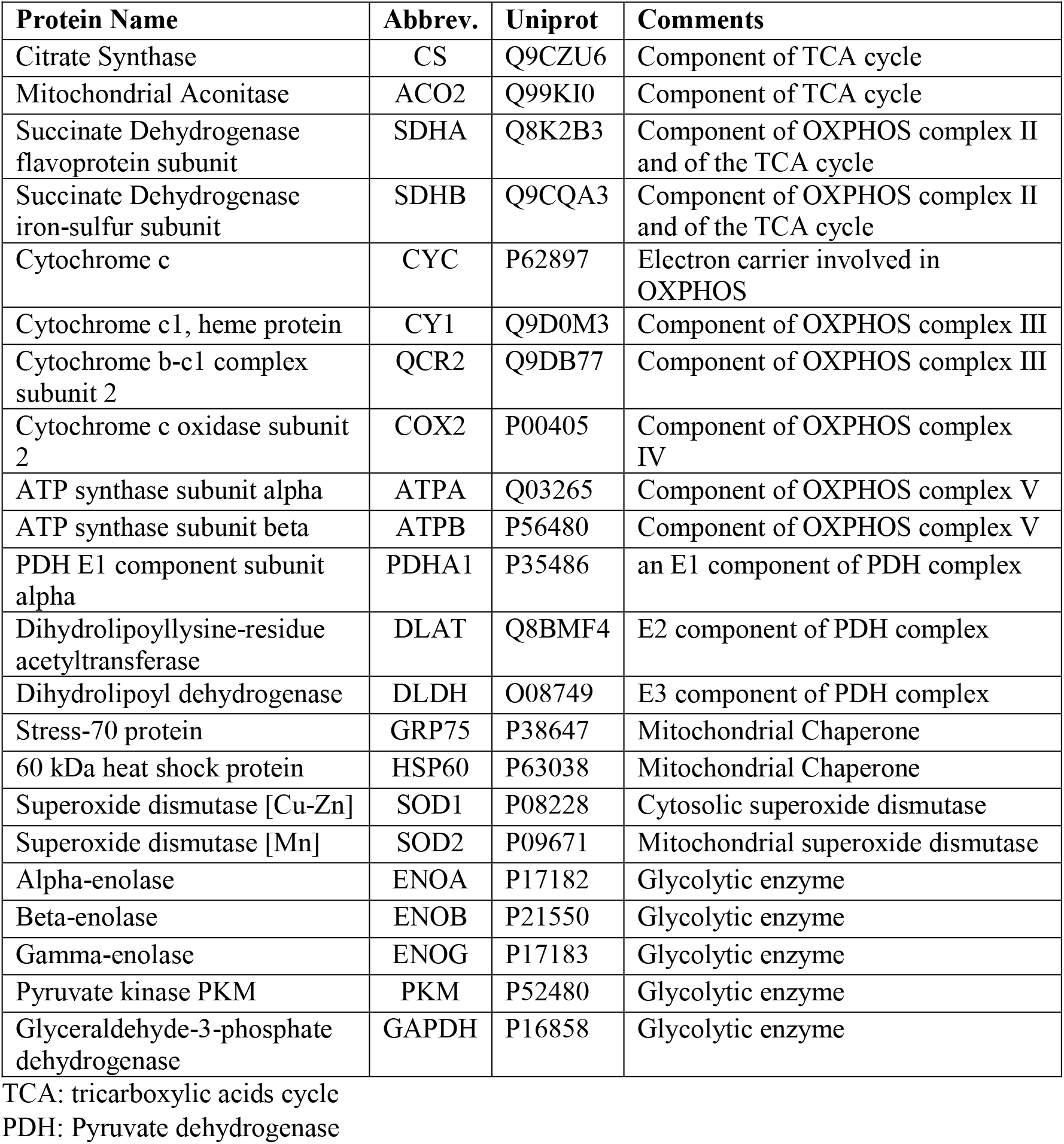
Proteins analyzed in the targeted proteomics analysis. The abbreviation used through this manuscript and the Uniprot accession code are indicated. Transitions used for detection of each protein are indicated in supplemental information.

**Figure 5.**
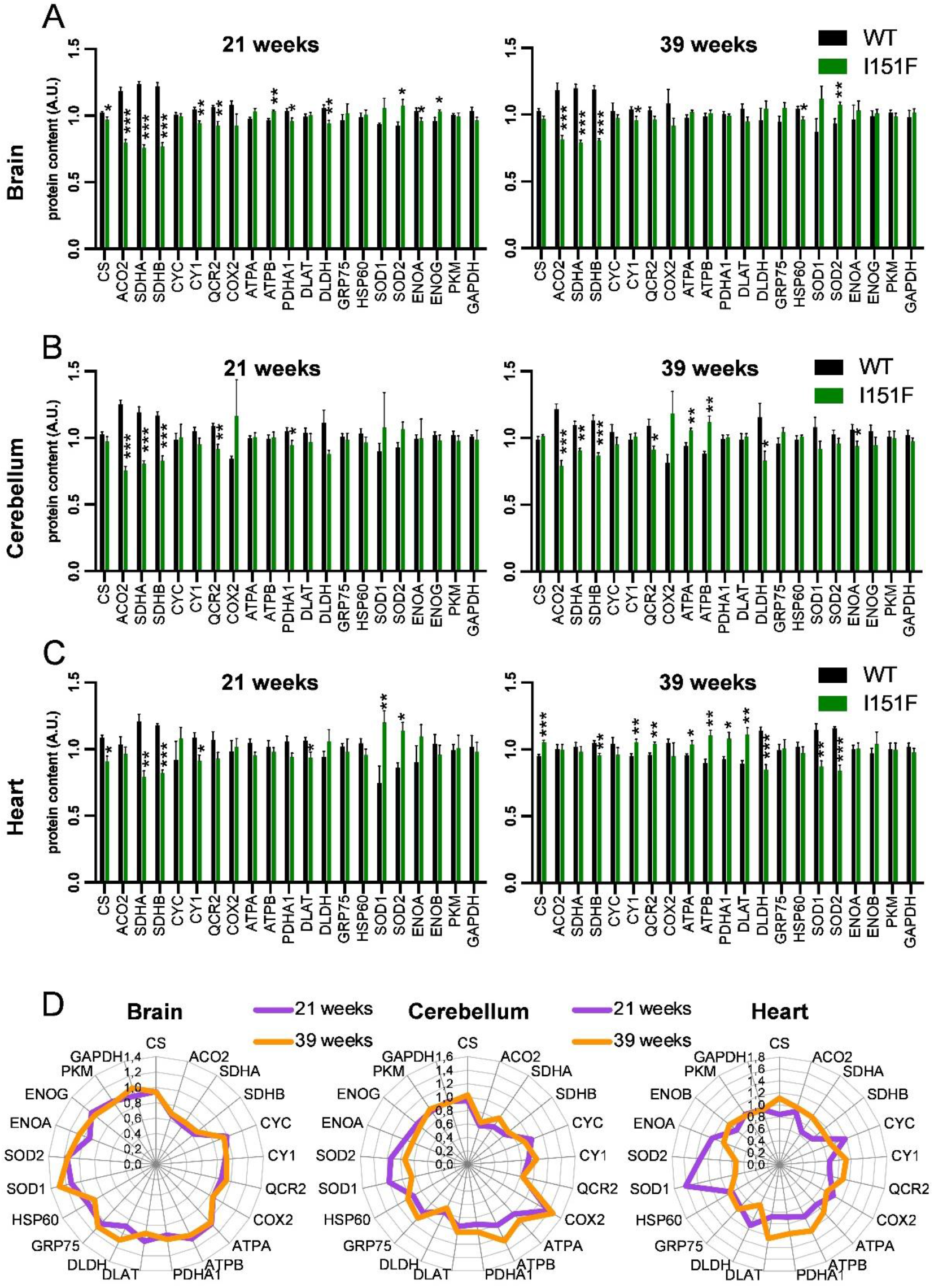
Analysis of the content of frataxin-related proteins by targeted proteomics. A to C, histograms show the relative content (in arbitrary units) of the measured proteins by targeted proteomics in brain, cerebellum and heart from 21 and 39-week old WT and Fxn^I151F^ mice. Data are represented as means±SEM from 4 (21-weeks) or 6 (39-weeks) mice. Significant differences p-values < 0.05(*), 0.01(**) or 0.001(** *) between WT and Fxn^I151F^ mice are indicated. D, net-representations indicate the relative changes observed the content of the indicated proteins in 21-weeks (violet) and 39-weeks (orange) old Fxn^I151F^ mice (relative to WT).

### Fxn^I151F^ mice present lower aconitase activity

ACO2 is a mitochondrial enzyme involved in the tricarboxylic cycle, where it converts citrate to isocitrate. It contains a 4Fe-4S cluster which is required for its enzymatic activity. As it is one of the most abundant iron-sulfur containing proteins, this protein has been commonly used as an indicator of the status of iron-sulfur centers in the cell. Therefore, we analyzed its content by western blot and also we measured total aconitase activity in brain, cerebellum and heart homogenates from 21-week and 39-week old mice. ACO2 is expected to account for most of aconitase activity in these tissues, as it represents between 80 and 90 % of all aconitase content according to PaxDb database. Western blot analysis confirmed the results observed in the targeted proteomic analysis, as the results obtained were similar to those obtained in the previous section: a marked decrease was observed in nervous system, while in heart no significant differences were observed between WT and Fxn^I151F^ mice (Figure 6). Regarding activity, we measured the ratio between aconitase and citrate synthase activities (ACO/CS ratio), which was used as a control for non-iron-sulfur mitochondrial enzyme. We observed significant decreased ACO/CS activity ratio in the three tested tissues, with more marked decreases in brain and cerebellum than in heart (Figure 6). Overall, the results obtained confirm that aconitase content and activity is most affected in the nervous system than in heart. They also suggest that decreased aconitase activity is mostly due to decreased ACO2 protein content, although the presence of some inactivated protein cannot be excluded (as in some of the analyzed condition, loss of activity is slightly higher than loss in protein content).

**Figure 6.**
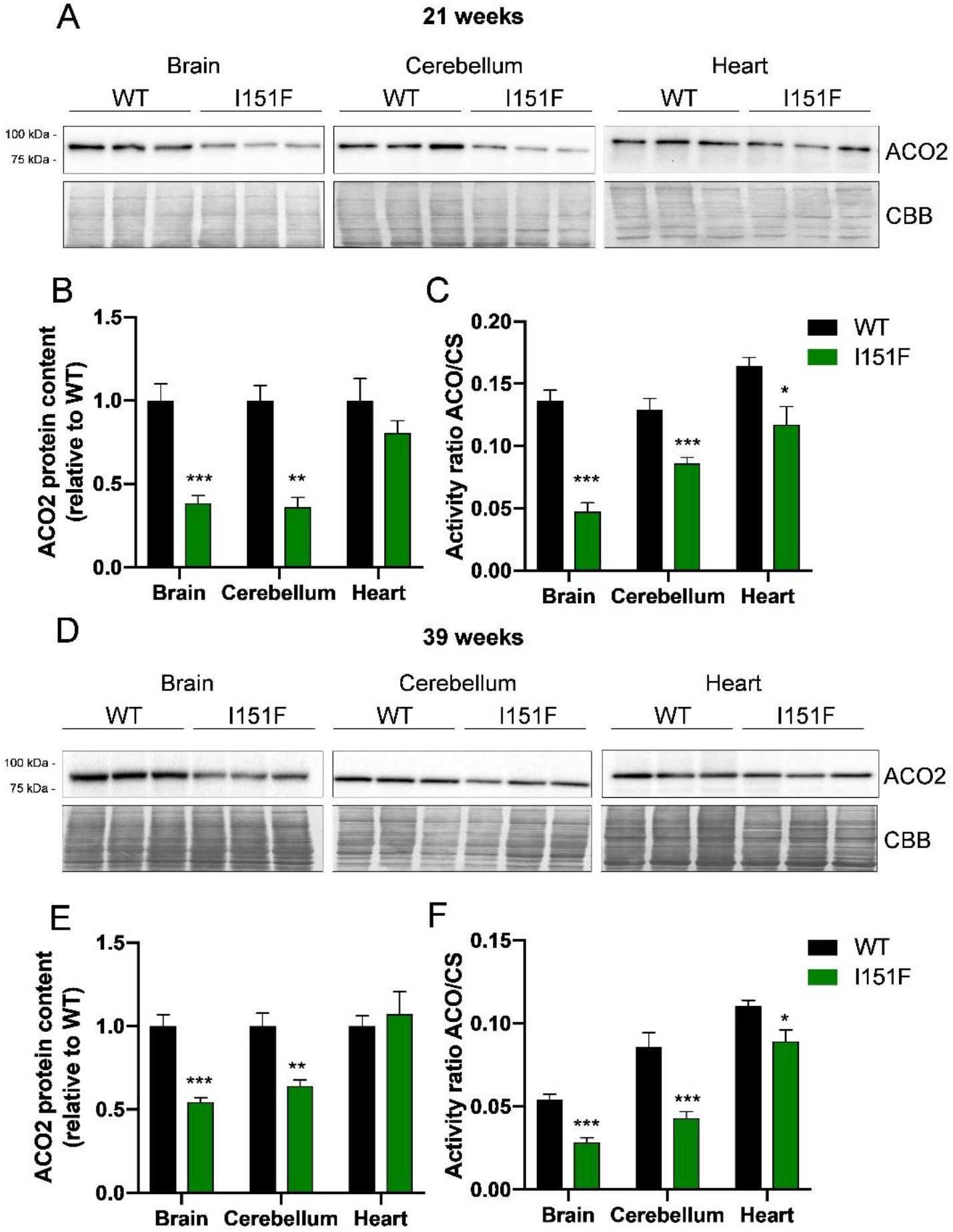
Aconitase content and activity in WT and Fxn^I151F^ mice. A, representative ACO2 western blot images from 21 week-old WT and Fxn^I151F^ mice. Brain, cerebellum and heart homogenates from three different mice were loaded on gels. B, ACO2 content in 21 week-old WT and Fxn^I151F^ mice. Histograms represent the means±SEM from at least 4 different mice. C, aconitase to citrate synthase activity ratio (ACO/CS) was measured in brain, cerebellum and heart homogenates from 21 week old WT and Fxn^I151F^ mice. Data represent the means±SEM from at least 5 different mice. D, representative ACO2 western blot images from 39 week-old WT and Fxn^I151F^ mice. Brain, cerebellum and heart homogenates from three different mice were loaded on gels. E, ACO2 content in 39 week-old WT and Fxn^I151F^ mice. Histograms represent the means±SEM from at least 4 different mice. F, aconitase to citrate synthase activity ratio (ACO/CS) was measured in brain, cerebellum and heart homogenates from 39 week-old WT and Fxn^I151F^ mice. Data represent the means±SEM from at least 5 different mice.

### Fxn^I151F^ mice present decreased content of OXPHOS complexes I and II

The targeted proteomics analysis also revealed changes in the content of several components of the OXPHOS system, notably complex II. In order to confirm these results with an alternative approach, we analyzed by western blot the content of 5 components of the complexes I to V (one protein per complex) in brain, cerebellum and heart from 21 and 39 week old mice. These proteins were NDUB8 from complex I, SDHB (complex II), QCR2 (complex III), COX1 (complex IV), and ATPA (complex V). None of the components of complex I had been included in the targeted proteomics analysis as they could not be properly quantified. The results obtained can are shown in figure 7. It can be observed that Fxn^I151F^ mice present a marked loss of NDUB8 from complex I and SDHA from complex II in all tissues analyzed (except heart at 39-weeks), and that this decreased content was already observed at 21 weeks of age. The results from SDHB are similar to that found in the targeted proteomics analysis, confirming the validity of the data obtained. Regarding complexes III to V, a small decrease in QCR2 content (from complex III) could be appreciated in cerebellum, as it had been previously observed in the targeted proteomics approach. No changes in ATPA content were observed by western-blot analysis. Overall, the results obtained by the targeted proteomics approach and the western blot analysis, indicate that Fxn^I151F^ mice present a deficiency in components of complex I and II, especially in the nervous system, while the components of the other complexes do not present major differences.

**Figure 7.**
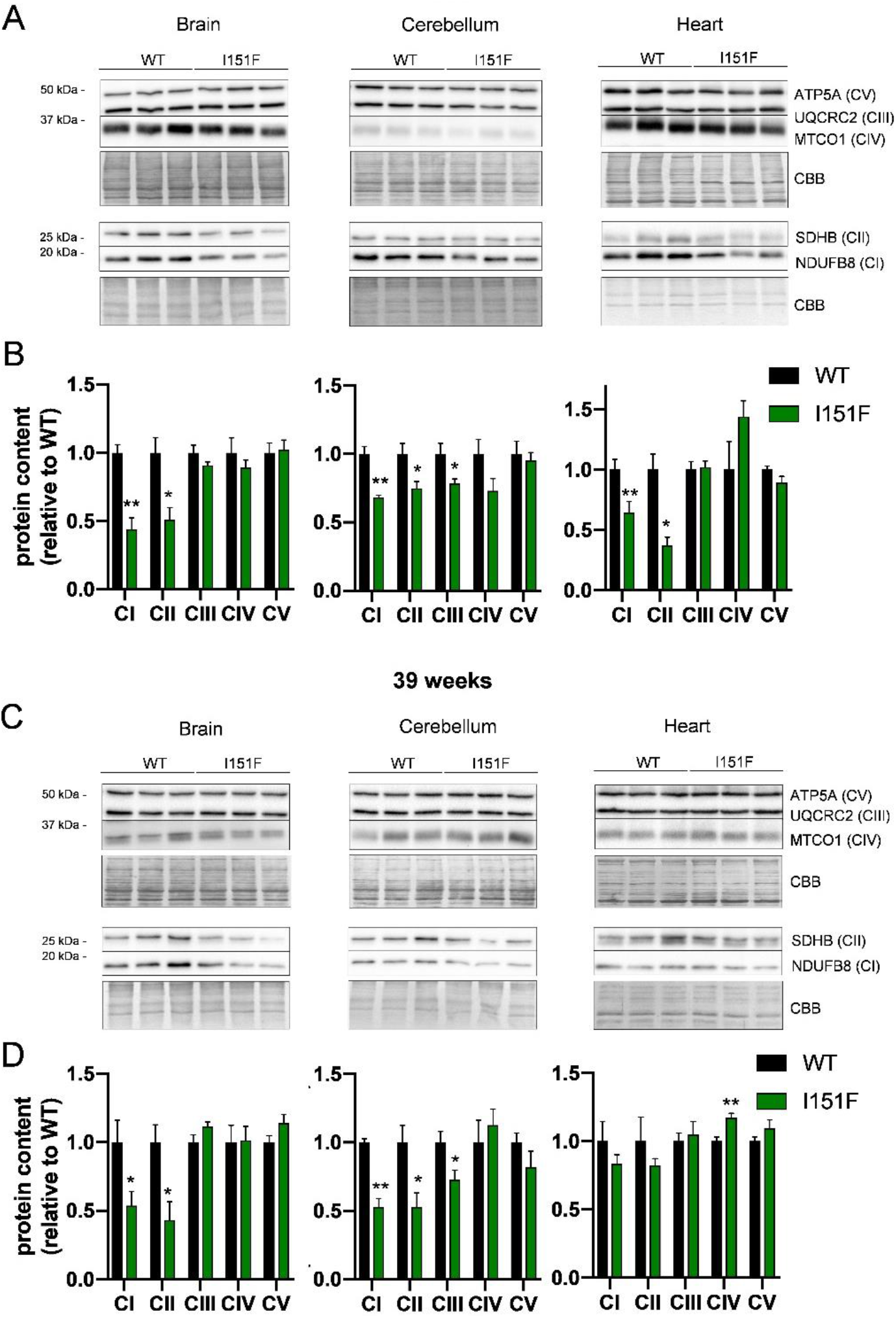
Analysis by western blot of components from OXPHOS system. The indicated components of the mitochondrial OXPHOS system were analyzed by western blot in brain, cerebellum and heart homogenates from 21 week and 39-week old WT and Fxn^I151F^ mice. A, representative western blot images from 21 week-old mice. Samples from three different mice were loaded on gels. B, content of the indicated proteins in 21 week-old mice. Histograms represent the means±SEM from at least 4 different mice. C, representative western blot images from 39 week-old mice. Samples from three different mice were loaded on gels. D, content of the indicated proteins in 39 week-old mice. Histograms represent the means±SEM from at least 4 different mice. Significant differences between WT and Fxn^I151F^ mice are indicated (p-values < 0.05(*), 0.01(**) or 0.001(** *)).

### Biosynthesis of lipoic acid is not compromised in Fxn^I151F^ mice

We also decided to analyze the content of protein-bound lipoic acid, a prosthetic group required for PDH activity. This analysis had two objectives: first, to complement the results from the targeted proteomics analysis, where several components of the PDH complex were analyzed; second, to further analyze the status of iron-sulfur clusters in Fxn^I151F^ mice, as synthesis of this cofactor requires lipoate synthase, which is an iron-sulfur containing enzyme. Western blot analysis of brain, cerebellum and heart lysates using antibodies raised against lipoate revealed the presence of two bands, at 70 and 50 kDa apparent molecular weight (Figure 8). The 70 kDa band (which is more intense) corresponds to lipoic acid bound to DLAT (the E2 component from PDH), while three lipoic acid containing proteins migrate at 50 kDa. These are the E2 component from alpha-ketoglutarate dehydrogenase (Dihydrolipoyllysine succinyltransferase, DLST), the E2 component from the branched chain alpha-ketodehydrogenase complex, and the PDH-binding component X (a structural subunit from the PDH complex) (Purroy *et al*, 2020). As indicated in figure 8, no significant differences where observed in DLAT-bound lipoic acid (70 kDa) between WT and Fxn^I151F^ mice. Decreased content in the 50kDa band was observed in 21-week old brain, but not in the other tissues analyzed. This band was also observed decreased in 39-week old mice, although in these mice differences did not reach statistically significance. Overall, these results indicate that protein-bound lipoic acid biosynthesis (which requires an iron-sulfur enzyme) is not compromised in Fxn^I151F^ mice.

**Figure 8.**
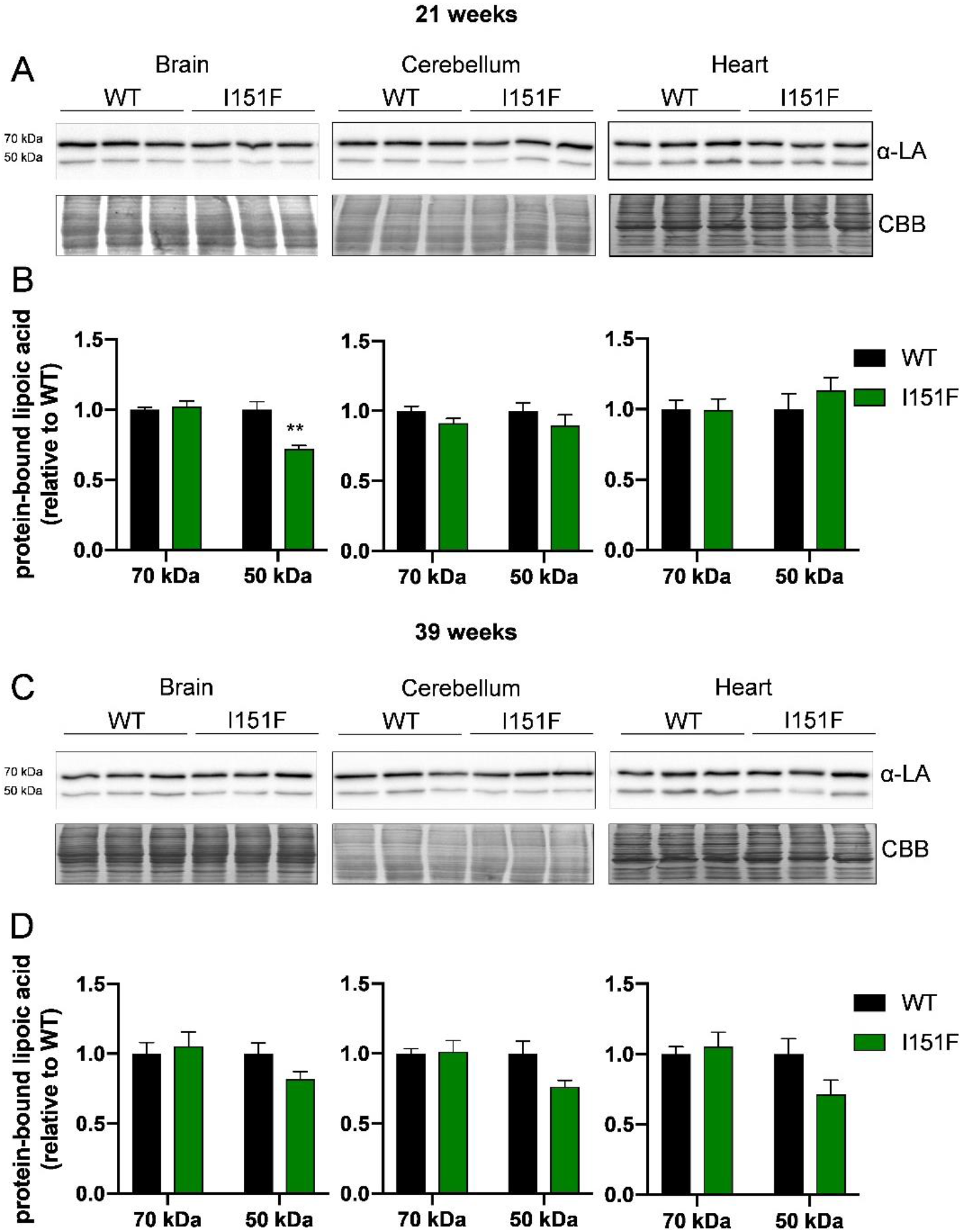
Analysis by western blot of protein-bound lipoic acid. Protein bound lipoic acid was analyzed using lipoic acid (LA) antibodies in brain, cerebellum and heart homogenates from 21 week and 39-week old WT and Fxn^I151F^ mice. The 70 kDa band corresponds to lipoic acid bound to DLAT, while three lipoic acid containing proteins migrate at 50 kDa (DLST, E2 from branched chain alpha-ketodehydrogenase complex, and PDH-X). A, representative western blot images from 21 week-old mice. Samples from three different mice were loaded on gels. B, intensity of the indicated bands in 21 week-old mice. Histograms represent the means±SEM from at least 4 different mice. C, representative western blot images from 39 week-old mice. Samples from three different mice were loaded on gels. D, intensity of the indicated bands in 39 week-old mice. Histograms represent the means±SEM from at least 4 different mice. Significant differences between WT and Fxn^I151F^ mice are indicated (p-value < 0.01(**)).

### HET mice do not present altered functional or biochemical parameters

Finally, in order to explore the effect of the I151F mutation in heterozygosis, we analyzed the performance of HET mice in an open field test, and also the content of some representative proteins in brain, cerebellum and heart. No significant differences were observed in the open field test, neither in velocity, distance travelled or number of crossings (Figure 9A). Regarding protein content, the analyzed proteins were CS, ACO2, SDHB, ATPB, SOD2, DLAT, GRP75 and PKM. This group includes representative proteins which experienced changes in the mutant mice (ACO2, SDHB, ATPB and SOD2) and also proteins which did not experience such changes but are representative of different pathways/functions. These proteins were analyzed in brain, cerebellum and heart from 39-week old mice by targeted proteomics. Results in figure 9B indicate that none of these proteins presented relevant changes in their content between HET and WT animals. A slight decrease in the amounts of CS and SDHB was observed in the cerebellum of HET mice, but these changes were lower than those observed in FXN^I151F^ mice and/or were not observed in brain and heart. Therefore, we can conclude that heterozygote mice do not present major biochemical nor functional alterations.

**Figure 9.**
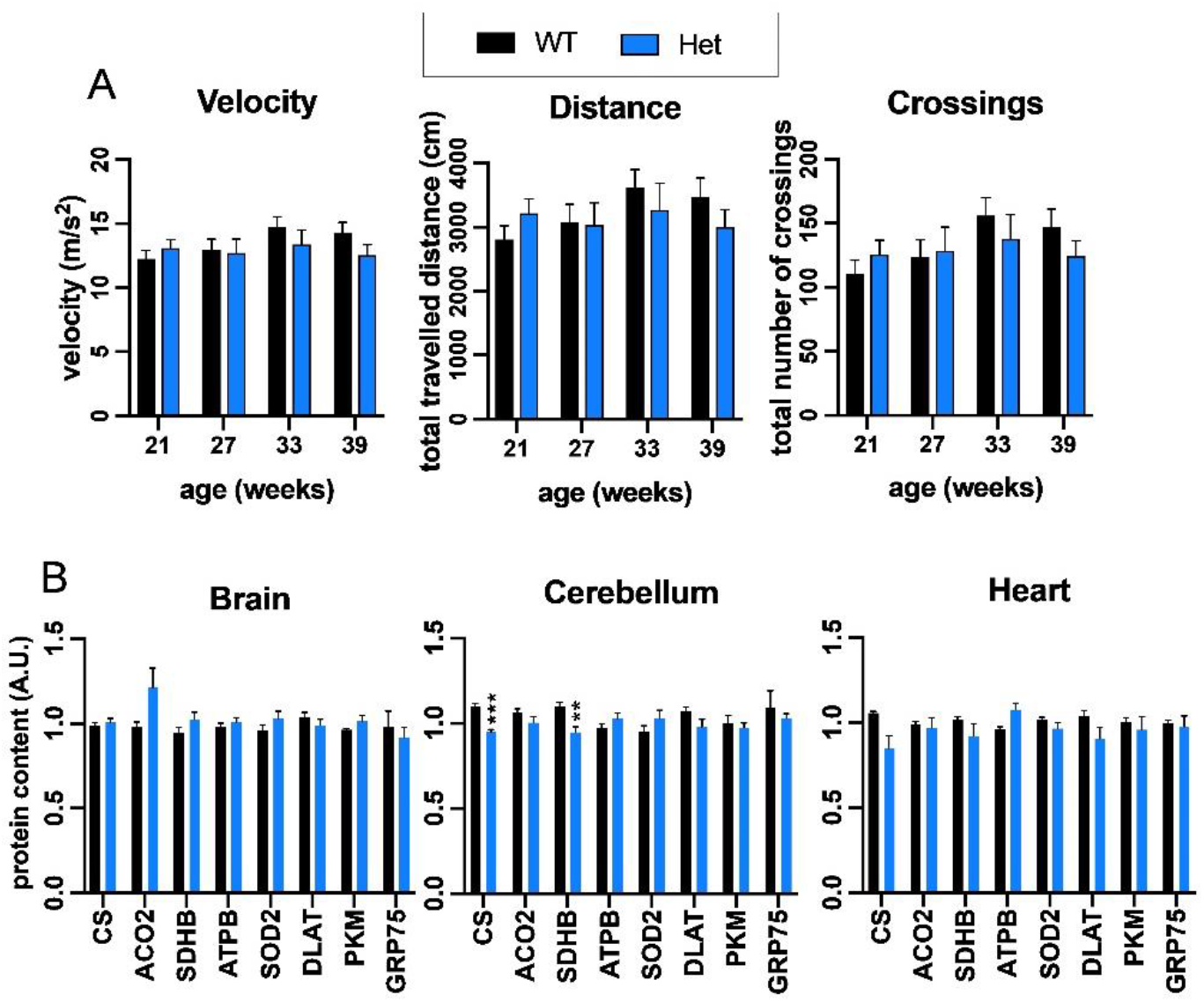
Analysis of HET mice. A, In an open field test, HET mice did not show significant decline in velocity, total distance traveled or number of crossings when compared with WT animals of the same age. Number of animals used in this analysis is indicated in supplemental table 1. B, Histograms show the relative content (in arbitrary units) of the measured proteins by targeted proteomics in brain, cerebellum and heart from 39-week old WT and HET mice. Data are represented as means±SEM from 4 mice. Significant differences between WT and HET mice are indicated (p-values < 0.05(*), 0.01(**) or 0.001(** *)).

## DISCUSSION

Several FA mouse models have been generated, which can be classified in four main categories: 1) tissue specific KOs, which present a total loss of frataxin in some tissues; 2) GAA-repeat mice models, which present a GAA expansion in the frataxin gene; 3) an inducible mutant, in which frataxin expression is repressed by a shRNA transgene placed under the control of a TET promoter (doxycycline-inducible) (Chandran *et al*, 2017); 4) point mutation models, in which mouse frataxin has been genetically modified by CRISPR to present a pathological point mutation. GAA-repeat mice models have the advantage of mimicking the most common mutation found in patients (the GAA expansion). Nevertheless, all the GAA-repeat models developed to date present a very mild phenotype, suggesting that the frataxin expression in these models is above (or slightly below) the pathological threshold required to trigger FA-like phenotypes in mice (reviewed in (Perdomini *et al*, 2013)(Ocana-Santero *et al*, 2021)). This limits its usefulness as a model to study the pathophysiology of the disease and also to test new therapeutic approaches, as no clear biomarkers of disease severity can be defined. Regarding models based on point mutations, a partial characterization of a G127V FXN mouse model (equivalent to G130V human pathological mutation) was recently published. This model has a severe phenotype, as mice harboring this mutation presented a substantially reduced number of offspring. Mouse embryonic fibroblasts derived from this model exhibit significantly reduced proliferation and bioenergetics alterations (Fil *et al*, 2020), but there is no published data about the consequences of this mutation in adult mice. In the present work we have performed a functional and a biochemical characterization of a FA mouse model based on the pathological mutation I154F. Therefore, this is the first FA mouse model based on a point mutation extensively characterized to date.

The clinical phenotype of patients carrying the I154F mutation (compound heterozygotes for the GAA triplet-repeat expansion and the point mutation) is similar from that of individuals homozygous for the GAA expansion (Filla *et al*, 1996), suggesting that mice carrying this mutation are a good model to study the pathophysiology of FA. Actually, the I154F mutation was reported to compromise the solubility of frataxin intermediate form and result in low levels of mature frataxin (Li *et al*, 2013). Therefore, the biochemical consequences of this mutation may be similar than those caused by the GAA expansion (as both cause decreased frataxin content). In this regard, in the present work we have observed that FXN^I151F^ homozygous mice present very low frataxin levels in all the tissues analyzed, confirming that the main consequence of this mutation is decreasing frataxin content. This marked loss of frataxin in all tissues is an advantage over other FA mice models, where frataxin deficiency is either mild or not present in all tissues. Our results also suggest that low residual frataxin content is enough to sustain viability in C57BL/6J mice. This observation is consistent with previous reports of mild phenotypes in GAA-repeat mice models. For instance, when 9 months old KIKO mice, which present a 70% reduction in frataxin content, were tested for 13 neurobehavioral tasks, significant differences were only observed in three of them (McMackin *et al*, 2017). In contrast, we have observed a more severe phenotype in the Fxn^I151F^ mice. First, they present decreased weight gain, which are observed from 14 weeks onward. Second, they present neurological deficits, which are manifested from week 22 onward. Third, they present marked biochemical alterations, which are observed before the appearance of the functional alterations (they are already observed in 21 week old mice).

The behavioral tasks performed indicate that Fxn^I151F^ mice exhibit several neurological deficits that are reminiscent of those observed in FA patients. Thus, Fxn^I151F^ mice displayed decreased locomotor activity, defects in their forelimb muscular strength, reduced motor coordination and balance and reduced hind and front limb stride length when compared with WT control. Also, the results obtained indicate that neurological defects are progressive, as they are not manifested until 22 weeks of age. Indeed, most of them are observed at older ages. In this regard, an interesting observation from this work is that biochemical alterations are observed before the onset of the functional disturbances, and that those alterations are not progressive in the nervous system (they are similar in 21 week old mice than in 39 week old mice). Among the proteins analyzed, we have found decreased content of NDUF8 (from complex I), SDHA and SDHB (from complex II), and ACO2. This observation is consistent with previous results in the KIKO mouse model, where complex I and II enzyme activities were also found decreased in cerebella from asymptomatic mice (Lin *et al*, 2017). NDUF8, SDHA, SDHB and ACO2 have in common that they either contain iron-sulfur clusters (SDHB and ACO2) or either belong to complexes containing them (complex I and II). Although, it could be hypothesized that these proteins are degraded due to the absence of iron-sulfur clusters, we should be cautious, since the Fxn^I151F^ mice do not have a general loss of iron-sulfur clusters. This is evidenced by the high aconitase content and activity found in heart, and also by the normal presence of protein-bound lipoate in most conditions analyzed (protein-bound lipoate requires an iron-sulfur containing protein for its biosynthesis (Ollagnier-de Choudens *et al*, 2000)). Therefore, the mechanism explaining the loss of these proteins may be more complex and could involve different regulatory pathways. It is worth reminding that iron-sulfur loss is not a universal characteristic of frataxin deficiency (Alsina *et al*, 2018a)(Bayot *et al*) and that studies in yeast indicate that iron-sulfur loss is caused by metabolic remodeling (Moreno-Cermeño *et al*, 2013; Alsina *et al*, 2018b).

The targeted proteomic analysis has also revealed changes in SOD1 and SOD2 protein content. These alterations are more marked in heart than in nervous system. Both enzymes are involved in superoxide scavenging and had been previously described either induced or decreased in frataxin-deficient cells or mice (Santos *et al*, 2001)(Sturm *et al*, 2004)(Chantrel-Groussard *et al*, 2001)(Irazusta *et al*, 2006). Interestingly, we have observed both phenomena, as both enzymes are induced in 21 week old mice, while their content is decreased in 39 week old mice. The mechanism causing this paradoxical behavior could be complex, as several transcription factors have been described to regulate superoxide dismutases expression. In this regard, the promoter of SOD1 gene contains binding sites for at least seven transcription factors and at least two of them, Nrf2 and NF-κB, could be responding to the oxidative stress conditions caused by frataxin deficiency (Milani *et al*, 2011). Moreover, it has also been described that NRF2 signaling may be inhibited in frataxin-deficent cells (Paupe *et al*, 2009)(Shan *et al*, 2013). Thus, SODs expression would depend on the balance between activation of these transcription factors by stress, and inhibition of NRF2 signaling. Such balance could be different at different ages, explaining the differences observed between 21 and 39 week old mice. It is worth commenting that a similar behavior was observed in hearts from the cardiac KO mouse model (MCK/frataxin), which showed a modest SOD2 induction between 2 and 7 weeks followed by a dramatic decrease at later ages (Seznec *et al*, 2005). Another potential explanation for the SODs decline observed in 39-week old hearts could be related to alterations in metal homeostasis caused by frataxin deficiency. In this regard,studies in yeast showed that activities of both SOD1 and SOD2 where compromised due to limited availability of Cu and Mn, which are their metal cofactors. Such limited availability of Cu and Mn would be caused by interferences related to disrupted iron homeostasis (Irazusta *et al*, 2010).

In this work we have also analyzed the content of three PDH containing components. In primary cultures of frataxin-deficient neonatal rat ventricular cardiomyocytes (NRVMs) we had previously observed decreased PDHA1 content and disturbances in the redox state of DLAT-bound lipoic acid (Purroy *et al*, 2020). These previous results indicated a 40% loss of PDHA1 content, and no major changes in DLAT or DLDH content. Indeed, treatment with the PDH activator dichloroacetate restored some of the defects found in frataxin-deficient NRVM (Purroy *et al*, 2020). The results obtained in the present work indicate a slight decrease of PDHA1 in 21-week old mice, while this protein was induced in hearts from 39-week old mice. Regarding DLDH, we have observed a 25 to 30% loss in its content in hearts and cerebellum from 39-week old Fxn^I151F^ mice. In this context, the previous results in rat cardiomyocytes indicated that DLAT-bound lipoic acid was more oxidized in frataxin-deficient cells, and the decreased content of DLDH in Fxn^I151F^ mice could cause a similar consequence in the mice. Therefore, we have observed alterations in the PDH complex, although the physiological consequences of these subtle differences, if any, would require further research.

In summary, the results obtained indicate that the FXN^I151F^ mice is an excellent model for FA research, as it presents low levels of frataxin content in all tissues and clear functional and biochemical biomarkers which resemble those observed in FA patients and models. Our results also indicate that frataxin deficiency impacts the OXPHOS system, aconitase activity and antioxidant defenses, both in the heart and the nervous system. Those biochemical alterations precede the appearance of neurological symptoms and present a different profile in brain, cerebellum or heart and. We can also conclude that the FXN^I151F^ mouse model is a very useful for investigating the pathological consequences of frataxin deficiency. Furthermore, the presence of clear biochemical and functional biomarkers indicate that this model can be used to test the effect of therapeutic approaches.

## MATERIALS AND METHODS

### Generation of FXN^I151F^ mice

The investigation with experimental animals conforms to the National Guidelines for the regulation of the use of experimental laboratory animals from the Generalitat de Catalunya and the Government of Spain (article 33.a 214/1997) and was evaluated and approved by the Experimental Animal Ethic Committee of the University of Lleida (CEEA). FXN^I151F^ heterozygous mice (C57BL/6J-F*xn*^*em10(T146T,I151F)Lutzy*^/J) were obtained from the Jackson Laboratory, Bar Harbor, ME USA (Stock Number 31922). To generate these HET mice, the CRISPR+guide targeted mutagenesis reagent was microinjected into C57BL/6J zygotes. A founder female was crossed with C57BL/6J to generate N1 HET mice which were also placed into mating to generate N2 HET mice. The mutant allele presented an ATC → TTC mutation in the I151 codon, which causes the expected I151F mutation. An additional silent mutation was present in T146 codon (ACC → ACT), while the remaining FXN coding sequence was free of errors. The silent T146T mutation was co-introduced with the I151F mutation as a PAM blocker. Intercrosses of HET animals were performed to generate the WT, HET and Fxn^I151F^ animals used in the present study. Genotyping of these mice was performed by sequencing a PCR product amplified from DNA extracted from tail biopsy specimens. The primers used were Fw: TTTCACACTTCCTGCCACCT; Rv: AGGCAGACAGCCGTAAAGTC. Animals were housed in standard ventilated cages at 12h light/dark cycle and fed with a normal chow diet ad libitum. Animals were weighed weekly. For isolation of tissues, animals were sacrificed by cervical dislocation at 21 or 39 weeks of age and dissected immediately. Isolated organs were snap-frozen in liquid nitrogen and stored at −80°C.

### Behavior analyses

The number of mice used for each analysis is summarized in supplemental table 1. The performed tests were Rotarod, Open Field, Hanging Wire and Paw Print. All tests were conducted by the same investigator and under dark light conditions. The number of mice used in each analysis is indicated in supplemental table 1. Mice were subjected to Rotarod every two weeks from 15 to 39 weeks of age. Open Field, Hanging Wire and Paw Print tests were performed at 21, 27, 33 and 39 weeks of age. The Rotarod test was performed in a LE 8200 Rotarod system (Panlab Harvard Apparatus). Animals were subjected to three trials in which, after 1 min at constant velocity (5 rpm), the rotarod was accelerated at 0.05 rpm/s. The latency to fall after acceleration was recorded and the average of the three trials was the used value. For Open Field test, a mouse was placed in the center of the field and its movements were recorded during 5 min using Smart Video Tracking software (v2.5.21, Panlab Harvard Apparatus). The measured parameters were total distance travelled, entries in zones (crossings) and average speed. The Hanging Wire test was performed to assess forelimb grip strength and consisted in testing the ability of the mouse to hang on a horizontally positioned wire for 30 sec. When the mouse fell from the wire, the latency to fall was recorded. Each mouse had three opportunities to achieve the 30 sec objective. The best mark of the three trials was the used value. Bedding material was placed underneath the wire to break the fall. For the Paw Print test, a runway of 55 cm long and 10 cm wide covered with laboratory paper was used. First, each mouse was placed into the runway during 30 s for habituation. Then, hind and fore paws were painted with pink and blue non-toxic paint respectively, and the mouse was placed at the beginning of the runway and allowed to walk. The papers were scanned and the stride length of the animals was measured using ImageJ software.

### Preparation of tissue homogenates

A piece of tissue sample (20–100 mg) was cut into 2–3 mm^2^ pieces. All the pieces were placed in 1.5 ml screw cap polypropylene tubes in the presence of lysis buffer consisting in 50 mM tris(hydroxymethyl)aminomethane (Tris) HCl pH 7.5 containing a protease inhibitor cocktail (Roche). For each 100 mg of tissue, 375 μl of lysis buffer were used. Glass beads (0.5–1.0 mm) were added to the mixture which was homogenized in a BioSpec Mini-Beadbeater. Following homogenization, SDS was added to the mixture at 4% final concentration. This homogenate was vortexed for 1 minute, heated at 98°C for 5 minutes, sonicated and subsequently centrifuged at 12000g for 10 min. Protein content in the supernatant was quantified using the BCA assay (Thermo Scientific).

### Protein quantitation by targeted proteomics

Tissue homogenates (50 μg of protein) were precipitated with cold acetone (9 volumes) and resuspended in 1% sodium deoxycholate, 50 mM ammonium bicarbonate. Then, proteins were subjected to reduction by 12 mM DTT and alquilation by 40 mM IAM. Mass spectrometry grade trypsin (SOLu-Trypsin, Sigma) was added to a final enzyme:substrate ratio of 1:50. After overnight digestion at 37°C, formic acid was added to precipitate sodium deoxycholate. The resulting peptide mix was purified and enriched using 100 μl Pierce C18 ZipTips. Eluted fraction from the C18 ZipTip was evaporated using a Concentrator Plus (Eppendorf) and peptides were resuspended in 3% acetonitrile plus 0,1% formic acid containing a heavy peptide standards mixture. Heavy peptides were obtained from JPT (SpikeTidesTM_L). All peptide samples were analyzed on a triple quadrupole spectrometer (Agilent 6420) equipped with an electrospray ion source. Chromatographic separations of peptides were performed on an Agilent 1200 LC system using a Supelco Bioshell A160 Peptide C18 column (1 mm x 15 cm). Peptides (up to 15 micrograms of protein digest) were separated with a linear gradient of acetonitrile/water, containing 0.1% formic acid, at a flow rate of 75 μl/min. A gradient from 3 to 60% acetonitrile in 45 minutes was used. The mass spectrometer was operated in multiple reaction monitoring mode. Transitions were obtained from SRM atlas or Prosit (Gessulat *et al*, 2019) and imported into Skyline software (MacLean *et al*, 2010), which was also used to analyze results. Once validated and optimized, the SRM assays were used to quantify all the analyzed peptides using scheduled SRM mode in a single run (retention time window, 120 s; cycle time, 1 sec). For calculating protein content, the light to heavy ratio of each peptide in each sample was first divided by the mean value of that peptide between all samples. Then, this value was normalized to the average value of all the proteins analyzed in each sample. The peptides and transitions analysed can be found in supplemental table 2.

### Western blot

After SDS-polyacrylamide gel electrophoresis, proteins were transferred to PVDF (Millipore, IPVH00010) or Nitrocellulose (Sigma_Aldrich, 10600093) membranes and blocked with I-block (ThermoFisher, T2015). The membranes were probed with the following primary antibodies: Frataxin (AbCam, ab219414), aconitase 2 (Sigma, HPA001097), lipoic acid (Calbiochem, 437695) and OxPhos (Invitrogen, 458099). Detection was performed using peroxidase conjugated secondary antibodies. Image acquisition was performed in a ChemiDoc MP system from Bio-Rad. Membranes were stained with Coomassie brilliant blue or Ponceau for normalization. When required, data was analysed by ImageLab software (Bio-Rad).

### Enzyme activities

A piece of tissue sample was cut into 2–3 mm^2^ pieces. All pieces where placed into tubes containing non-denaturing lysis buffer consisting of Tris-HCl 50 mM at pH 7.4, protease inhibitor cocktail (Roche) and sodium citrate 2.5 mM. Glass beads (0.5–1.0 mm) were added and tissues were homogenized using a BioSpec Mini-Beadbeater. Then, Triton X-100 was added at a final concentration of 0.5%. Homogenized tissues were centrifuged at 13000 rpm at 4°C during 5 min and the supernatants were placed into new tubes. Aconitase activity was measured in Tris-HCl 50 mM at pH 7.4, containing 1 mM of sodium citrate, 0.2 mM of NADP, 0.6 mM of manganese chloride and 0.25 units of isocitrate dehydrogenase (Sigma-Aldrich, I2002). NADPH formation was measured at 340 nm during 120 s. Citrate synthase activity was measured with a coupled assay to reduce 5,5′-dithiobis-(2-nitrobenzoic acid) (DTNB). Briefly, tissue extracts were added to Tris-HCl 100 mM pH 8.1 with 0.4 mg/ml of DTNB and 10 mg/ml of Acetyl-CoA. Absorbance was measured at 412 nm during 120 s. Then, 8.5 mg/ml of oxaloacetate were added into the cuvette and the absorbance was measured again at 412 nm during 120 s for the detection of reduced DTNB. Values are presented as a ratio of aconitase activity *versus* citrate synthase activity.

### Statistical analysis

For statistical analysis, the two-tailed Student’s t-test was used to assess the significance of the differences between data. The p-values lower than 0.05(*), 0.01(**) or 0.001(***) were considered significant.

## Acknowledgments

The authors would like to thank Aamir Zubery and Cathleen Lutz from the Rare and Orphan Disease Center at the Jackson Laboratory for Fxn^I151F^ HET mice model generation. This model was generated thanks to the BeHeard Challenge hosted by the Rare Genomics Institute (CA, USA). This work has also been funded by grants from Association Française de L’Ataxie de Friedreich (AFAF) and from Ministerio de Economía y Empresa (MINECO, Spain, SAF2017-83883-R). We thank Roser Pané for technical assistance, as well as the Proteomics and the Animal services from Universitat de Lleida, and the metabolomic services from IRBLLeida.

## Author contributions

MMC and ASA performed most of the experiments and analyzed the data. EB and FD assisted in the collection and preparation of mouse tissues. EC assisted in the biochemical analyses. All authors provided technical support and suggestions for the project and for the manuscript. JR and JT conceived the project and supervised the study. MMC and JT designed the experiments, analyzed and interpreted data, and wrote the manuscript.

